# Integrated spatial morpho-transcriptomics predicts functional traits in pancreatic cancer

**DOI:** 10.1101/2025.03.12.642933

**Authors:** Dennis Gong, Rachel Liu, Yi Cui, Michael Rhodes, Jung Woo Bae, Joseph M. Beechem, William L. Hwang

**Affiliations:** Massachusetts Institute of Technology, Division of Health Sciences and Technology, Cambridge, MA; Center for Systems Biology, Department of Radiation Oncology, and Krantz Family Center for Cancer Research, Harvard Medical School and Massachusetts General Hospital, Boston, MA; Broad Institute of MIT and Harvard, Cambridge, MA; Bruker Technologies, Inc. Seattle, WA

## Abstract

Analyses of patient-derived cell lines have greatly enhanced discovery of molecular biomarkers and therapeutic targets. However, characterization of cellular morphological properties is limited. We studied cell morphologies of human pancreatic adenocarcinoma (PDAC) cell lines and their associations with drug sensitivity, gene expression, and functional properties. By integrating live cell and spatial mRNA imaging, we identified KRAS inhibitor–induced morphological changes specific for drug-resistant cells that correlated with gene expression changes. We then categorized a large panel of patient-derived PDAC cell lines into morphological (e.g., polygonal, irregular, spheroid) and organizational (e.g., tightly aggregated, multilayered, dispersed) subtypes and found differences in gene expression, therapeutic targeting potential, and metastatic proclivity. In human PDAC tissues, we identified prognostic expression signatures associated with distinct cancer cell organization patterns. In summary, we highlight the potential of cell morphological information in rapid, cost-effective assays to aid precision oncology efforts leveraging patient-derived in vitro models and tissues.

## Introduction

The size, shape, and organization of cells in culture represents a fundamental characteristic that distinguishes cellular identity, state, and function (*1*). Thousands of cell line models have been derived over the past century (*2*), each having distinct structural and organizational features that correspond to phenotype and tissue lineage (*3*). Cancer cell lines have previously been described to exist on a spectrum between epithelial-like and mesenchymal shapes (*4*, *5*). More aggressive and invasive cancer cells often exhibit mesenchymal morphology (*6*, *7*), a hallmark of the epithelial-mesenchymal transition (EMT) (*4*). Still, a broad degree of morphologic and organizational variation is not adequately captured by this simple binary schema.

More detailed characterization of cell morphology can provide a high-dimensional “fingerprint” associated with behaviors such as movement, response to perturbation, and death. Diverse patterns of motility, blebbing, cell adhesions, axon-like projections, nuclear morphology, and lipid metabolites can all be visualized by phase imaging (*8*). Cancer cells can change their morphology in response to drug treatment, which may highlight targetable drug resistance mechanisms. For instance, cells treated with DNA damaging agents begin to express DNA damage response proteins and undergo senescence to stop replication (*9*), taking on a distinctive flat and irregular morphology with increased cell area. Genetic alterations can also lead to changes in cell morphology. For example, mutations in genes regulating the cytoskeleton (e.g. Rho family GTPases) can alter cell shape and motility (*10*) and cell morphology can even predict mutational status (*7*). Microenvironmental ligands such as TGF-beta (*11*) as well as culturing conditions including mechanical substrate and ECM composition (*12*, *13*) also have potent morphological remodeling effects.

Large-scale image-based screens have recently enabled comprehensive analyses of the effects of various perturbations on cell morphology in single cell lines (*14–16*). However, the baseline heterogeneity in morphology across different cell lines remains largely unquantified, presenting an opportunity to integrate these measurements with gene expression profiling and dependency resources like the Cancer Cell Line Encyclopedia (CCLE) (*17*, *18*). Additionally, the shared and distinct morphological changes in response to perturbations across diverse cell lines are not well understood. An alternative screening strategy—assessing the remodeling effects of a few perturbations across many cell lines—has not yet been conducted at scale (*19*). Such an approach could reveal novel associations between morphology and -omics data, such as dependency and gene expression, and how these relationships shift under perturbation.

To address this gap in knowledge, we studied a large panel of human PDAC cell lines in response to clinically-deployed compounds including KRAS inhibitors (e.g. MRTX1133, RMC-6236) and chemotherapy (e.g. 5-FU, gemcitabine). We pioneered a new single-cell framework, Spatial Morphology and RNA Transcript Analysis (SMART), to integrate cell morphology and transcriptomic state using a combination of holotomography (Nanolive 3D Cell Explorer 96focus), CellPainting (*20*), and spatial molecular imaging (SMI; Bruker/Nanostring CosMx) (*21*). Utilizing SMART, we provide evidence of distinct cytoskeletal adaptations in response to KRAS inhibition and chemotherapy that correlate with drug sensitivity.

More broadly, to holistically evaluate the translational significance of cell morphology in patient-derived model systems, we performed analysis of -omics data from the Cancer Dependency Map (*22*) (DepMap), functional experiments including clonogenicity and invasion assays, and the CellPainting assay to quantify and categorize the morphological and organizational states present in PDAC cell lines. We identified molecular features of cells associated with their respective morphologies, finding that genetic dependency is predictive of morphology using an XGBoost (*23*) machine learning model. Using our categorization schema, we performed functional assays to discover morphologies and organizational patterns associated with invasion, stem-like features, and metastasis. Finally, we identified correlates to tissue morphology and identified strong associations between basal transcriptional subtype, which is associated with worse prognosis and treatment resistance (*24*), and small cell clusters, as well as classical transcriptional subtype and large clusters.

Altogether, we provide a robust framework for integrating morphological profiling into -omics workflows, offering a powerful approach to uncover novel biomarkers, characterize drug responses, and elucidate mechanisms of therapeutic resistance.

## Results

### Spatial Morphology and RNA Transcript (SMART) Analysis to Characterize Treatment Induced Remodeling of Cellular Morphology and Transcriptomic State

Cell morphology is a dynamic and cell state specific biomarker that can be investigated at different length scales and by visualizing different molecular and structural features. To comprehensively capture morphological phenotypes and their molecular correlates, we developed a multimodal framework utilizing multiple independent assays, which we termed Spatial Morphology and RNA Transcript (SMART) analysis (**Figure 1A**). First, we pioneered an ultrahigh (200 nm) resolution live cell phase imaging technology called holotomography (Nanolive 3D Cell Explorer 96focus) to measure cell shape dynamics in response to perturbations such as drug treatment. Additionally, we employed a CellPainting assay that provided higher throughput characterization and quantification of structural features such as actin fibers. To directly correlate morphology and associated transcriptional features, we integrated spatial molecular imaging (SMI; Bruker/Nanostring CosMx), which enables high-plex spatial transcriptomics in addition to static imaging data to link morphology and transcriptome. Each of our SMART measurements provide single cell data, overcoming both inter- and intra-cell line morphologic heterogeneity. Furthermore, in contrast to dissociative single cell approaches, which perturb cell morphology and transcriptomic state, our *in situ* SMI method directly enables accurate cell shape and transcriptome measurements in a single assay, while preserving cell-cell interactions that may influence cell state.

**Figure 1.**
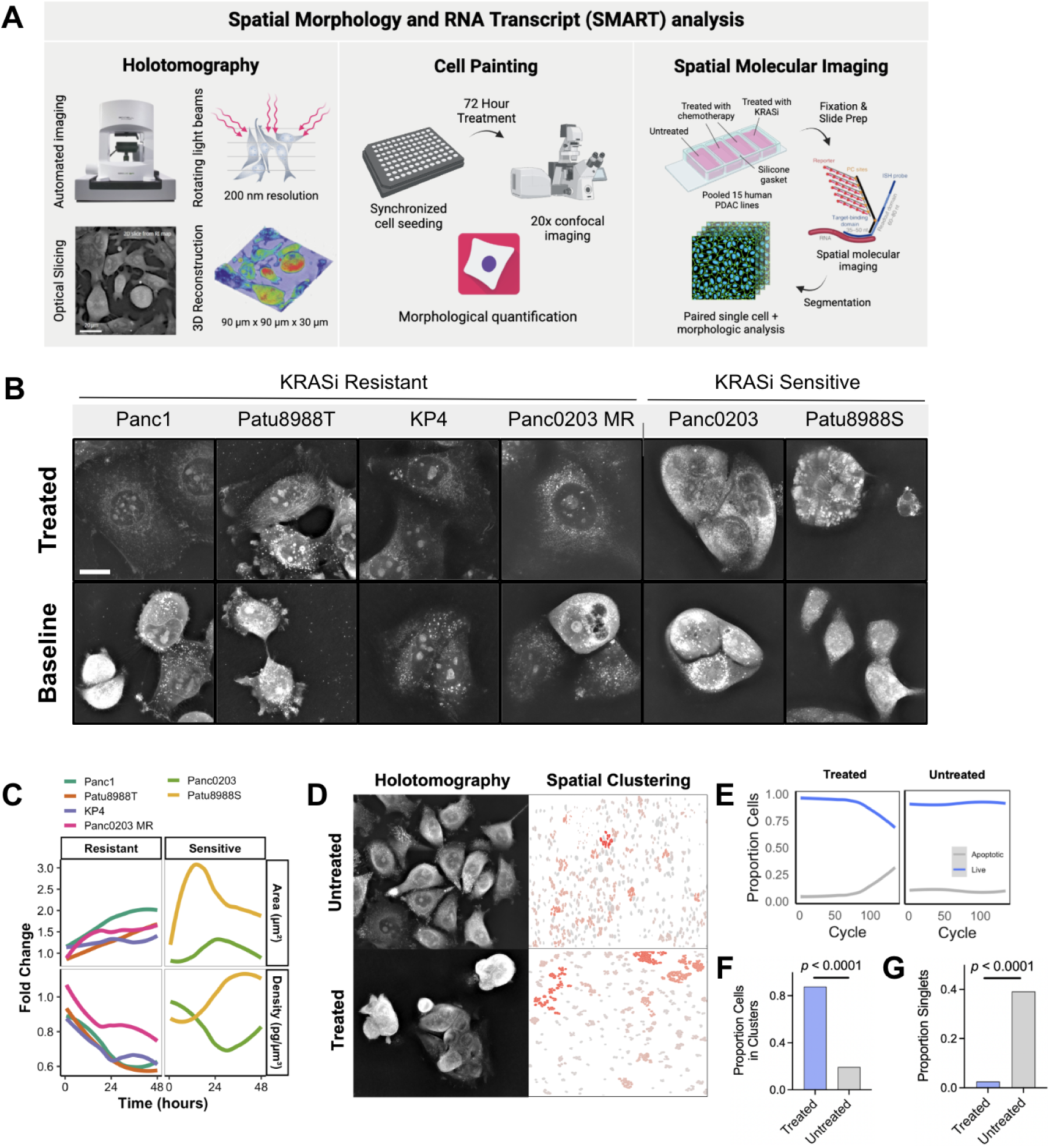
Dynamic morphologic and organizational remodeling in PDAC cell line models in response to KRAS inhibition by phase holotomography. (A) Schematic of Spatial Morphology and RNA Transcript (SMART) analysis framework. (B) Representative images of cell lines profiled with phase holotomography following 48 hours of KRAS inhibitor (RMC-6236) treatment and at baseline conditions. Scale bar = 20 µm. Panc0203 MR refers to the Panc0203 parental cell line grown to resistance to MRTX1133 in increasing drug concentration (*37*). (C) Quantification of cell object area and density throughout the treatment course with RMC-6236, stratified by RMC-6236 sensitivity. (D) Holotomography and clustering analysis of an additional KRAS inhibitor sensitive cell line, AsPC1, in KRAS inhibitor treated and untreated conditions. (E) Quantification of cell death of AsPC1 cells in treated and untreated conditions. Cycle length is 20 minutes. (F-G) Quantification of AsPC1 cells in clusters versus singlets in KRAS inhibitor treated and untreated conditions.

We applied SMART to study the treatment response of PDAC cell lines to KRAS inhibitors, due to their emerging clinical relevance in this highly treatment refractory disease. More than 90-95% of pancreatic adenocarcinomas harbor *KRAS* gain of function mutations (*25*), which have recently become targetable using both allele specific and pan-KRAS mutation small molecule inhibitors (*26*, *27*). Specifically, we chose RMC-6236, a RAS(ON) multi-selective inhibitor of KRAS currently in Phase II trials (*26*); and MRTX1133, an allele specific KRAS G12D inhibitor also in Phase II trials (*28*) for our studies. We hypothesized that changes in morphology may serve as a predictive biomarker for early cellular response to a drug perturbation including potential resistance mechanisms.

First, we performed live cell holotomography to visualize morphologic and organizational changes in patient-derived PDAC cell lines (**Figure 1B**). In a panel of six cell lines, four of which are resistant (AUC > 0.65) and two of which are sensitive (AUC < 0.50) to KRAS inhibition, we identified distinct morphologic changes associated with treatment sensitivity. Resistant cell lines enlarged their cell area without increasing cell mass, while sensitive cell lines clustered together forming tightly aggregated clumps that reduced in size over time, likely due to cell death (**Figure 1B,C**). To validate our findings, we took another sensitive cell line, AsPC1, and treated it with the KRAS^G12D^ specific inhibitor MRTX1133 prior to performing live cell holotomography (**Figure 1D** *left*). Similar to the sensitive cell lines Patu8988S and Panc0203, we observed some cell death following 24 hours of treatment (**Figure 1E**), but most notably, there was induction of tight clustering in response to KRAS inhibition (**Figure 1D**).

To further investigate the causes of treatment associated-morphologic changes, we harnessed another SMART assay, SMI, to make concurrent spatially-resolved morphology and transcriptomic measurements, which can be integrated with the dynamic morphological changes captured by holotomography (**Figure 1**). We performed SMI on AsPC1 cells treated with MRTX1133 to quantify clustering over a larger imaging window and also to identify gene expression changes in clusters versus singlets (**Figure 1D** *right*). Globally, there were significantly more cells in clusters of five or more touching cells in the treated versus untreated condition, and significantly less singlets, defined as cells without any touching neighbors (**Figure 1F,G**). Spatially-resolved gene expression analysis identified an increase in cell junction gene expression including various integrins (e.g. *ITGB1*, *ITGA3*, *ITGA6*) in cell clusters relative to singlets and in treated conditions relative to untreated conditions.

Given the heterogeneity in treatment response across resistant and sensitive cell lines, we sought to characterize and quantify treatment induced morphological changes at a broader scale. We first assembled a panel of 15 PDAC cell lines with a range of treatment sensitivities to RMC-6236 and MRTX1133. We then performed modified CellPainting (**Methods**) to directly visualize the morphology of individual cells after 72 hours of drug treatment. In addition to KRAS inhibitors, we included additional compounds with clinical relevance to PDAC as controls, including 5-FU, a thymidylate synthase inhibitor that is the backbone of the widely used FOLFIRINOX multi-agent chemotherapy regimen (*29*, *30*) and gemcitabine, a nucleoside analog, used as part of an alternate chemotherapy regimen (*31*). We performed confocal fluorescence imaging with a 20x objective in 15 of our cell lines in both the untreated (DMSO carrier control) and drug treated settings, totaling 75 conditions in arrayed format. Using CellProfiler (*32*), we segmented 14,488 cells and quantified cell morphology, organization, and staining intensity features for downstream analyses (**Figure S1A**).

We performed principal component analysis (PCA) and calculated a uniform manifold approximation and projection (UMAP) on normalized morphological measurements. Measurements related to cell size as well as compactness/eccentricity, a measure of deviation from a perfectly circular shape, were represented as top features in the first two principal components (**Figure S1B-D**). Cell lines did not cluster separately, indicating a significant degree of shape heterogeneity within each cell line (**Figure 2A**).

**Figure 2.**
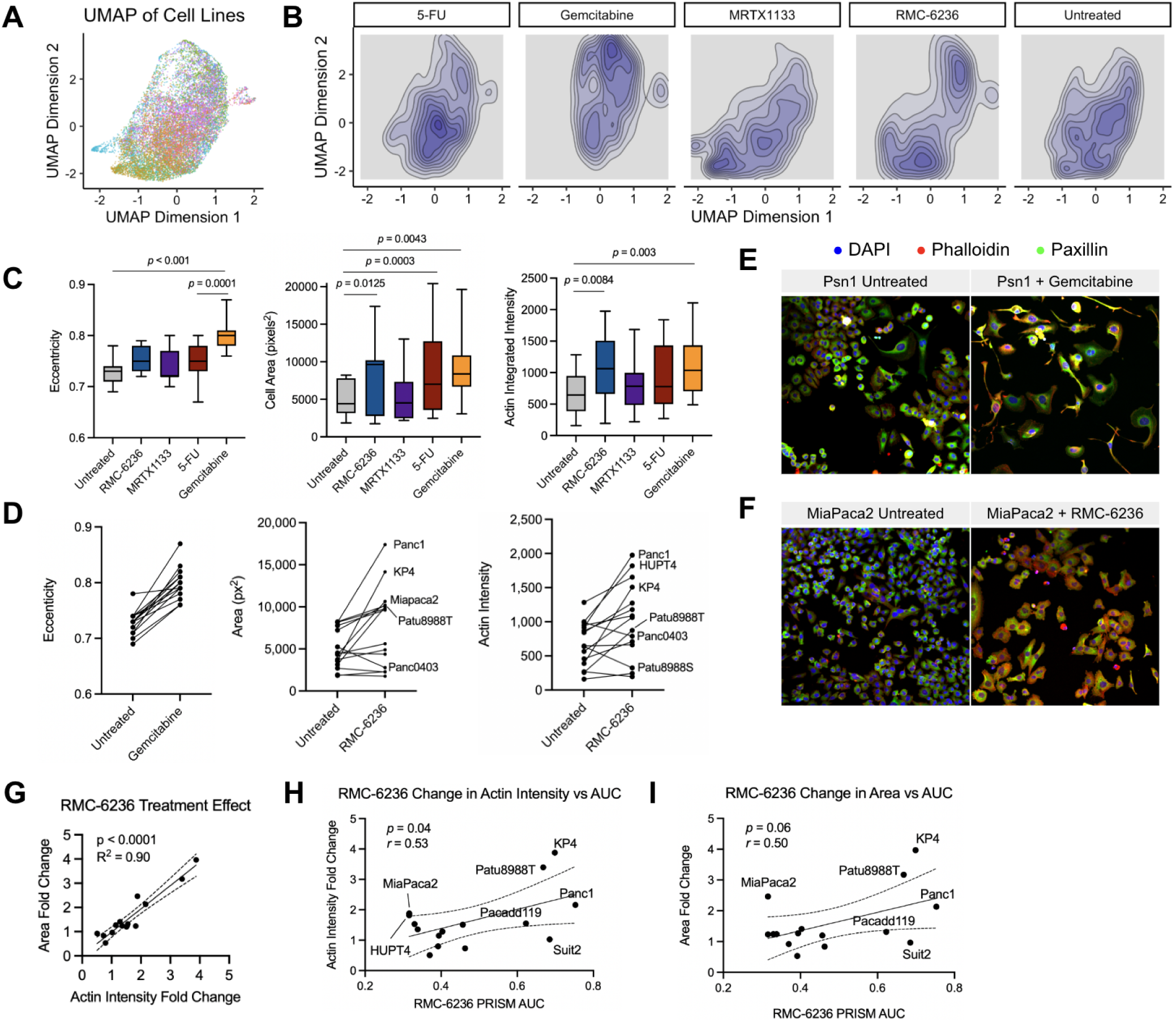
Treatment associated remodeling of cell morphology by CellPainting. (A) UMAP projection of individual cells across all conditions (untreated, gemcitabine, 5FU, MRTX1133, and RMC-6236 treated) pseudo colored by cell line. UMAP is clustered based on CellProfiler calculated morphological features. (B) Density plot of cells in different treatment conditions on the UMAP projection in (A). (C) Boxplot of calculated eccentricity, cell area, and actin integrated intensity features stratified by treatment. (D) Change in eccentricity, area, and actin intensity measurements for each cell line. (E) Representative image of Psn1 cells treated with gemcitabine or carrier control. (F) Representative image of MIA PaCa2 cells treated with RMC-6236 or carrier control. (G) Correlation of actin intensity and cell area fold change when treated with RMC-6236 across the panel of cell lines. Each dot represents an average of measurements across all profiled cells within each cell line. (H, I) Correlation of change in actin intensity and cell area (y-axis) with RMC-6236 drug sensitivity (x-axis) measured in PRISM pooled viability assay (DepMap).

There was global treatment induced remodeling of morphology in all treatment conditions, especially within the cell area and eccentricity measurements (**Figure 2B,C**). Of the two KRAS inhibitors, RMC-6236 treatment exhibited the more significant morphological shifts with highly correlated changes in cell area and actin intensity (**Figure 2C-E,G**). However, these shifts were heterogeneous across cell lines, whereas gemcitabine uniformly caused morphological changes in eccentricity in all cell lines tested (**Figure 2D,F**). Certain cell lines including Panc1, KP-4, MIA PaCa2, and Patu8988T exhibited a marked increase in cell area upon treatment with RMC-6236 while others such as AsPC1, Patu8988S, and Panc0403 showed no change or even featured decreased cell area in response to KRAS inhibition, supporting our phase holotomography findings (**Figure 1B**). Interestingly, these patterns of morphological change showed a correlation with sensitivity to RMC-6236 in a pooled PRISM assay (**Figure 2H,I**), suggesting that treatment resistance can manifest as morphologic changes. Our results indicate that cytoskeletal changes are a critical indicator of response and sensitivity to treatment and motivate additional experiments to determine if targeting cytoskeletal remodeling via focal adhesion kinases has synergy with KRAS inhibition.

Gemcitabine elicited the largest change in eccentricity relative to the untreated condition, suggestive of senescent morphology driven by cell cycle arrest. This was characterized by increased cell area, increased abundance of cell projections, reduced cell-cell contacts, and increased actin staining intensity (**Figure 2C,F**). These changes also occurred in the 5-FU treated cells, indicating a potential class effect of chemotherapy, but the magnitude of changes was not as significant as with gemcitabine at the IC50 for each drug. Senescence associated morphological changes have previously been observed in PDAC cell lines and DNA damaging agents such as gemcitabine are at least additive with senolytic agents in the preclinical setting (*33*).

### Morpho-Transcriptomic Measurements of Pooled Cell Lines

Our next goal was to link morphological shifts to transcriptional content across our entire panel of cell lines. To achieve this scale in a cost effective manner, we applied our SMI assay on pools of cells encompassing all 15 of our profiled PDAC cell lines in each of our previously defined treatment conditions (**Methods**). For Panc1 cells which we used to optimize our assay, we were able to detect nearly 700 unique genes and a mean of nearly 8,000 transcripts per cell using a 1000-plex RNA imaging panel (**Figure S2D**). We tested slide coating with Poly-L-lysine to improve cell adherence. Coated slides generally had better cell adherence and nearly equivalent performance metrics (**Figure S2D,E**), enabling us to analyze approximately five-fold more cells per assay. In our pooled assay, we were able to achieve a mean of 429 genes and 5,482 transcripts per cell after filtering, and similarly strong reproducibility across technical replicates (**Figure S2F,G**). The overall lower genes and counts per cell in the pooled assay is likely secondary to relative differences in cell size and transcript content across cell lines (**Figure S2H**). Importantly, these metrics were consistent across treatment conditions as measured by correlation of mean transcript counts per gene across technical replicates (**Figure S2I**).

To analyze this high dimensional dataset, we first performed Leiden clustering and visualized a UMAP projection with distinct clusters (**Figure 3A**) (*34*, *35*). We developed a similarity metric based on ridge regression to map bulk RNAseq profiles from DepMap onto clusters to determine whether clusters separate by cell lines (**Methods, Figure S3A**). Most clusters cleanly mapped to a single cell line (**Figure S3B**). For example, the group of cell clusters labeled 1, 2, 24, and 21 all cleanly mapped to Suit2, and clusters 3 and 6 cleanly mapped to MIA PaCa2. However, some clusters mapped to multiple cell lines, making definitive annotation more difficult. We ultimately assigned clusters based on their optimal mapping and calculated the composition of the cell line mixture across each treatment condition (**Figure 3B**). Reassuringly, we identified expansion or contraction of cell populations that was consistent with their known sensitivity to certain compounds. For example, RMC-6236 resistant cell lines KP-4, Patu8988T, and Panc1 all expanded relative to other cells in the RMC-6236 condition relative to untreated, while sensitive lines including MIA PaCa2, Patu8988S, and Psn1 all contracted (**Figure 3B**).

**Figure 3.**
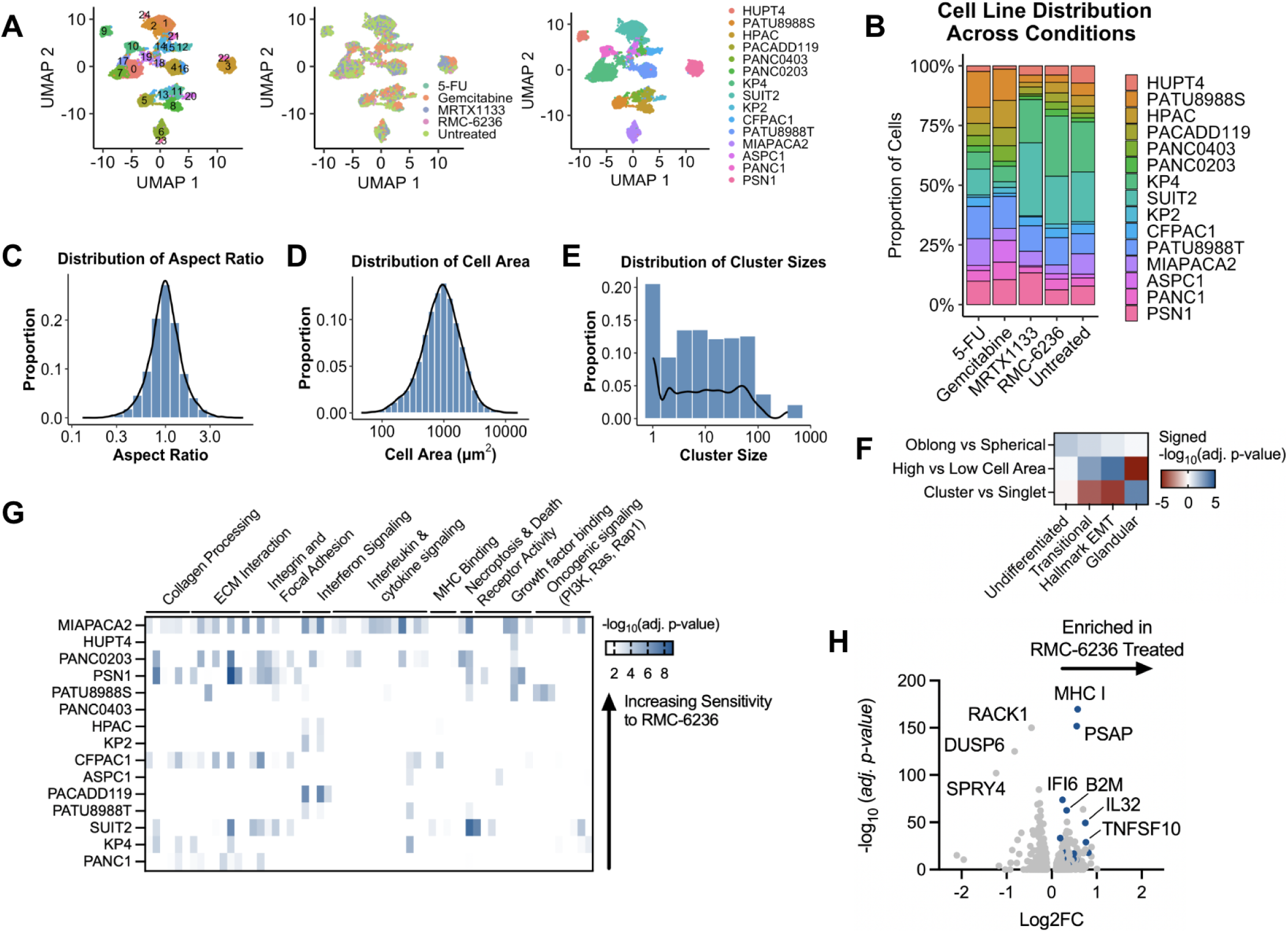
Correlation of morphology and transcriptional state at single cell resolution. (A) UMAP projection of single cell transcriptome profiles labeled by cluster assignment (left), treatment condition (middle), and cell line annotation (right). (B) Stacked barplot depicting cell line distributions across treatment conditions. Bars are represented as proportions and are colored according to cell line. (C-E) Histograms and probability distribution functions (black lines) of the aspect ratio (C), cell area (D), and cluster sizes (E) as defined by DBSCAN (**Methods**). (F) Heatmap depicting enrichment score (color) of cell state signatures (*36*) in each morphologic comparison (rows). (G) Clustered heatmap of enriched gene sets (columns) in RMC-6236 treated vs untreated conditions in each cell line (rows). Color density correlates with enrichment magnitude. (H) Volcano plot of enriched genes in RMC-6236 treated condition across all cell lines. Highlighted blue colored dots are related to inflammatory signaling mentioned in text.

In addition to annotating cell lines, we also sought to stratify cells by their morphological features. We defined three feature sets: cell area, cell aspect ratio, and cell aggregation (**Figure S3C**). We plotted distributions of these variables (**Figure 3C-E**, **S3D-E**) and stratified cells into groups. We compared cells with aspect ratio greater than 1.5:1 (termed oblong; 26.4% of total cells) with those exhibiting aspect ratios of 1.2:1 or below (termed spherical; 40.7% of cells). For cell area, we divided cells by bottom (<575 µm^2^) or top (>1440 µm^2^) quartile. Finally, we stratified cell aggregation using the spatial clustering algorithm DBSCAN (**Methods**). Cells with minimal contact with neighboring cells (singlets; 20.6% of total cells) were compared to cells that aggregated into clumps of 5 or more cells (56.5% of total cells).

Notably, distinct expression patterns of cell state signatures, including morphobiotypes (*36*) derived from *in situ* microdissected human PDAC specimens, were observed across these strata. Oblong shape was associated with a signature derived from microdissected PDAC cells with undifferentiated appearance (NES: 1.40, *p_adj._* = 0.024). Cells with large areas were enriched for EMT signatures including a signature derived from microdissected PDAC cells with structures transitioning away from ductal morphology (NES: 1.58, *p_adj._* = 3.18e-4) and the Hallmark EMT gene set (NES: 1.68, *p_adj._* = 1.35e-5), and depleted for glandular morphology signatures (NES: -1.96, *p_adj._* = 1.58e-10). The association of EMT signatures with enlarged cell area is consistent with our finding that cells resistant to KRAS inhibition increase their cell area. EMT signatures were previously found to be strongly enriched in mouse models of PDAC treated to resistance with MRTX1133 (*37*). Finally, cells in clusters were enriched for glandular signatures (NES: 1.40, *p_adj._* = 0.024) and depleted for EMT signatures (Hallmark EMT NES: -1.81, *p_adj._* = 4.24e-6; Transitional Morphobiotype NES: -1.73, *p_adj._* = 7.98e-5) (**Figure 3F**).

To study treatment effects on transcriptional and morphological features, we performed GSEA on the top differentially expressed genes between the treated and untreated cells in each condition (**Figure S4A-D**). Of note, we observed several KEGG terms enriched in the RMC-6236 treated condition, including focal adhesion (*p_adj._*= 1.68e-6), PI3K-AKT signaling (*p_adj._* =6.25e-5), and ECM-receptor interaction (*p_adj._* = 3.32e-4). Interestingly, PI3K focal copy number gains have been observed in mouse tumors treated with tool compound RMC-7977, an analog of RMC-6236 (*38*). KRAS withdrawal models as well as KRAS mutated PDAC lines with primary resistance to KRAS inhibitors have also been reported to utilize ECM derived focal adhesion signaling as an acute resistance mechanism (*39*, *40*).

In response to chemotherapy, we observed increased expression of inflammatory factors (**Figure S4A,B**). This finding is consistent with several known chemoresistance mechanisms including extracellular release of damage associated molecular patterns (DAMPs), which trigger secretion of interleukins and chemokines (*41*), and the senescence-associated secretion of inflammatory cytokines, chemokines, and growth factors (*42*). Cell adhesion related gene sets (e.g. GO:BP regulation of cell-cell adhesion) were enriched across all treatment groups relative to untreated cells. The cell adhesion response has commonly been reported as an ECM-mediated drug resistance mechanism to multiple classes of drugs (*43*, *44*).

Morphologically, we observed that all pooled cell lines increased their cell area in response to gemcitabine (**Figure S4E**), similar to our arrayed CellPainting experiments (**Figure 2**, **S4F-H**). However, certain shifts in morphology, especially in KRAS inhibitor treated conditions, occurred in a cell line dependent manner also similar to observations in the arrayed CellPainting assay (**Figure 2**, **S4F-H**). To investigate heterogeneity in the transcriptional response to KRAS inhibition, we performed differential expression analysis in a cell line specific manner. Using the top differentially expressed genes in the treated versus untreated condition for each cell line, we performed GSEA and clustered all the enriched terms. We visualized the clusters as a heatmap (**Figure 3G**) to identify shared and cell type specific changes in gene expression. Several modules emerged including gene sets relating to collagen processing, the extracellular matrix, integrin and focal adhesion signaling, immune response signatures for cytokine, interleukin, and interferons, as well as oncogenic growth signaling pathways (e.g. PI3K, Ras, Rap1) (**Figure 3G**).

Consistent with the known immunostimulatory effect of KRAS inhibition (*45*), MHC I expression was significantly increased in all cell lines (**Figure 3H**). Other differentially expressed genes enriched in KRAS inhibitor treated cells included *PSAP*, which encodes a secreted glycoprotein previously reported to reduce lymphocyte infiltration (*46*); *TNFSF10*, which encodes TRAIL, another secreted molecule capable of inducing cancer cell apoptosis; *B2M* which encodes an MHC I subunit; and *CD74* which encodes a portion of the MHC II complex (**Figure 3H**). Interferon response genes including *IFI6*, *ISG15*, *ITM2B*, *IFIT1*, *IFIT3*, and *IFITM1* were also significantly enriched in a subset of cell lines. In particular, MIA PaCa-2, which is sensitive to RMC-6236, but had a marked increase in cell size when treated with RMC-6236 (**Figure 2E**), had amongst the highest enrichment of interferon gene signature expression as measured by spatial molecular imaging of interferon gene transcripts. Thus, our pooled SMART assay recovered both shared and cell line specific transcriptional and morphologic responses to KRAS inhibition and chemotherapy.

### Morphologic and Organizational Classification of Human PDAC Models

Our next goal was to holistically evaluate the translational significance of cell morphology in patient-derived cell line avatars as a potential biomarker. We reasoned that it is essential to first define the landscape of morphologic diversity across patient-derived models. In addition to our cohort of 15 human PDAC cell lines used for treatment induced morphologic analysis, we identified 33 additional cell lines previously profiled in DepMap with publicly available phase-contrast images (*18*, *47*, *48*). These patient-derived cell lines encompass a broad range of age, sex, tumor site, and genetic diversity and thus provided a diverse cohort for morphologic, transcriptomic and functional characterization in this study.

We categorized cell lines by their predominant morphological appearance (i.e. polygonal, irregular, spheroid) and organizational pattern with neighboring cells (i.e. aggregated, multi-layered, dispersed) (**Figure 4A**, **S5**). Polygonal cells had well-formed edges typically associated with aggregation into distinct clumps of cells with close contact among neighbors, but without physical contact with other clumps. Irregular cells featured non-symmetrical shapes, typically with a spindle-like appearance and/or projections emanating from the cell body. Finally, spheroid morphology was characterized by spherical cells. There were also distinct organizational patterns. The aggregated pattern was characterized by close contacts among cells in a cluster, such that discernible gaps were not visible. The multi-layered pattern was characterized by cells growing on top of each other in the z-plane. Finally, the dispersed pattern describes a configuration where the cells were more evenly distributed over the plated surface compared to the aggregated and multi-layered patterns.

**Figure 4.**
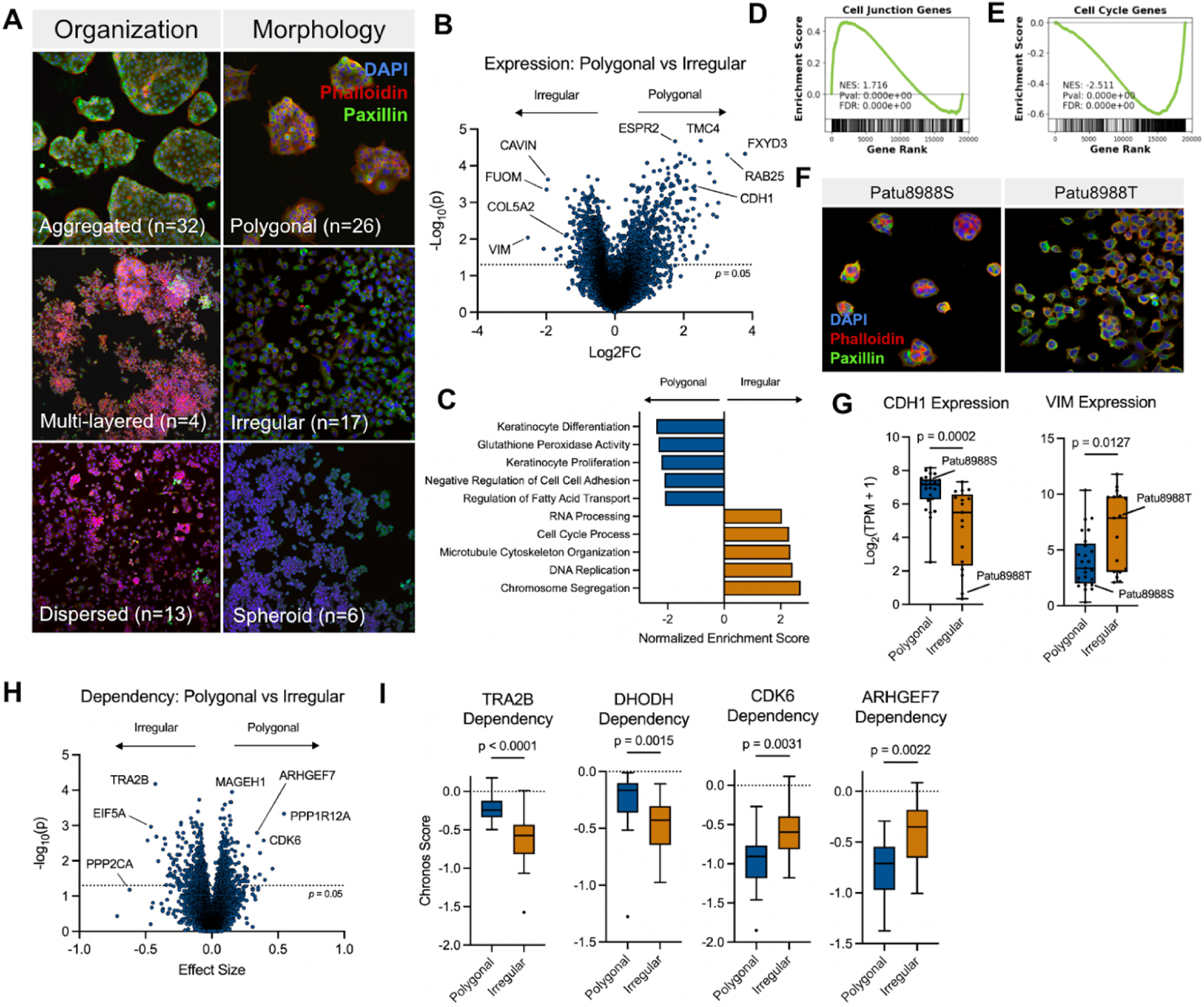
Transcriptional, and dependency correlates of morphology. (A). Categorization of cell morphologies (polygonal, irregular, and spheroid) and organizational patterns (aggregated, multi-layered, and dispersed) of cell lines in our study. Categories were assigned based on microscopic visualization in culture and image analysis (CellProfiler). (B) Volcano plot depicting differentially expressed genes between polygonal and irregular cell lines utilizing bulk RNA-seq data from the CCLE. (C) Barplot highlighting top 5 GSEA terms enriched in polygonal (left) and irregular (right) cell lines. (D) GSEA enrichment plot of GO:BP Cell junction organization in polygonal cell lines relative to irregular cell lines. (E) GSEA enrichment plot of GO:BP Cell cycle in polygonal cell lines relative to irregular cell lines. (F) Boxplot of *CDH1* and *VIM* transcript expression stratified by polygonal or irregular morphology. (G) CellPaint imaging of Patu8988S and Patu8988T. (H) Volcano plot depicting dependency differences between polygonal and irregular cell lines. Effect size (x-axis) is calculated as the difference in mean dependency. (I) Boxplots of notable dependency differences (*TRA2B*, *DHODH, CDK6 ARHGEF7*) stratified by morphology.

The aggregated organizational pattern was most closely aligned with the polygonal cell morphology, though certain cell lines with irregular and spheroid morphology exhibited heterogeneous organization (e.g. some cells aggregated, others dispersed). Morphologies were generally conserved across passages, freeze-thaw cycles, different standard media conditions (e.g. RPMI or DMEM-based complete media), and cell confluency. Notably, cellular organization was partially dependent on cell confluency, with ultra-low confluences supporting more dispersed organization and high confluency forcing some degree of aggregation in all cell lines. Thus, our organizational and morphologic classifications were performed at intermediate confluence during the log growth phase (i.e. 40-70% confluency).

### Gene Expression and Dependency Correlates of Morphology

We analyzed transcriptomic correlates of cell morphology by querying RNA expression data associated with each of the 49 PDAC cell lines in our combined cohort. We first focused on differences between polygonal and irregular morphology given the abundance of these morphologies in our cohort (n = 26 polygonal, n = 17 irregular). We performed differential expression analysis (**Figure 4B**) and gene set enrichment analysis of the differentially expressed genes (**Figure 4C-E**). Relative to irregular cells, polygonal cell lines were enriched for cell junction genes (normalized enrichment score/NES: 1.716, *p_adj._* < 0.0001), as well as gene sets for keratinocyte development and proliferation (GO:BP Keratinocyte differentiation, GO:BP Keratinocyte proliferation). In contrast, irregular cell lines were broadly enriched for cell cycling gene sets (NES: 2.511, *p_adj._* < 0.0001) including chromosome segregation (GO:BP Chromosome segregation, GO:BP Chromosome organization), DNA replication (GO:BP DNA Replication), and microtubule cytoskeletal organization (GO:BP Microtubule based process, GO:BP Microtubule cytoskeletal organization). While there were a limited number of cell lines with spheroid morphologies (n = 6), we noted that compared to both polygonal and irregular cell lines, they had the highest enrichment of cell junction and adhesion formation genes (GO:BP Cell adhesion, GO:BP Cell junction assembly), as well as neuron projection development (GO:BP Neuron projection development).

Polygonal cell lines were notably enriched for glutathione peroxidase activity, which protects against oxidative stress. GPX4, a key enzyme in this family, was previously identified as a selective dependency in mesenchymal cell lines due to their increased reliance on fatty acid metabolism (*49*, *50*). Despite these findings, the expression of *GPX4* and other key regulators of fatty acid transport including *CYP4F2*, *CYP4A11*, *ACSL1*, and *ACSL5* also have significantly reduced expression in more mesenchymal appearing irregular cells relative to polygonal cell lines. The increased dependency on GPX4 in mesenchymal cell lines, despite reduced expression of its encoding gene and related pathway components, suggests that mesenchymal cells are unable to buffer against reduction in GPX4 levels and thus have greater sensitivity to GPX4 inhibition (*51*).

Patu8988S and Patu8988T are two cell lines derived from the same PDAC liver metastasis that exhibit distinct morphological, functional, and transcriptional properties (**Figure 4F**). The Patu8988S cell line features polygonal morphology, aggregated organization, and high levels of *CDH1* consistent with its overall epithelial phenotype (*52*) (Mann-Whitney U test; *p* = 0.0002) (**Figure 4B,G**). In contrast, Patu8988T has irregular morphology and dispersed organization, expresses higher levels of *VIM* (Mann-Whitney U test; *p* = 0.0127), and exhibits a markedly higher proliferation rate, consistent with its overall mesenchymal phenotype (*53*) (**Figure 4B,G**). To ensure that none of the morphologies map to atypical transcriptomes that are not represented in patient tumors, we queried a dataset quantifying the similarity of cell line models to transcriptional profiles from resected tumors in the TCGA (*54*). The morphological and organizational patterns identified were each represented among cell lines representative of patient-derived tumors (**Figure S6**).

Next, we analyzed DepMap dependency data, quantifying differences in Chronos score between polygonal and irregular morphologies. We discovered distinct dependencies associated with morphologic state, including some therapeutic targets of interest in PDAC (**Figure 4H,I**). *CDK6*, the cell cycle kinase targeted by palbociclib and other CDK4/6 inhibitors used in hormone receptor-positive (HR+) breast cancers, had higher dependency in polygonal PDAC cell lines (Mann-Whitney U test; *p* = 0.0031). Supportively, HR+ breast cancers also most commonly exhibit a luminal epithelial morphology (*55*). Another polygonal specific dependency was *ARHGEF7* (Mann-Whitney U test; *p* = 0.0022), which encodes a RAC1 guanine nucleotide exchange factor that is involved in cytoskeletal remodeling. This gene had enriched dependency in PDAC, biliary tract, and oral squamous cell carcinomas, and was recently found to sensitize PDAC cell lines to CHK1 inhibition (*56*). *DHODH*, encoding dihydroorotate dehydrogenase in the pyrimidine biosynthesis pathway, is a dependency specific for the irregular morphology (Mann-Whitney U test; *p* = 0.0015). DHODH is a druggable metabolic dependency in subsets of PDAC cell lines that are resistant to *KRAS* ablation (*57*).

The most significant differential dependency between polygonal and irregular morphologies in our cohort was the highly conserved RNA splicing factor *TRA2B*, which exhibits a mean difference in Chronos dependency score of 0.427 in irregular versus polygonal lines (Mann-Whitney U test; *p* < 0.0001). TRA2B has critical functions in development, including for somitogenesis (*58*), cortical neurodevelopment (*59*), and spermatogenesis (*60*), and has been reported to contribute to oncogenesis in mesenchymal cancers including osteosarcoma (*61*) and in laryngeal squamous cell carcinoma models (*62*). Interestingly, *TRA2B* was expressed significantly higher in primary (*p* = 0.0001) and metastatic (*p* = 0.038) PDAC relative to physiologic pancreas tissue and was associated with worse overall survival in a cohort of 178 PDAC patients (*p* = 0.032) (**Figure S7A,B**).

Finally, we applied a tree-based gradient boosted machine learning algorithm (XGBoost) to measure the predictive value of expression or dependency data for classifying polygonal or irregular morphology. Using a leave-one-out cross validation approach we found that transcript expression measurements were a weaker predictor of morphology (AUC = 0.701) than dependency (AUC = 0.828) (**Figure S7C,D)**. When the two feature sets were combined into a single model, accuracy for predicting morphology (AUC = 0.845) was only marginally improved over the dependency model alone (**Figure S7D,E**). *TRA2B* dependency was by far the most predictive feature in each of the models utilizing dependency as quantified by improvement in accuracy (gain) when added to the model (**Figure S7D,E**).

### Functional Correlates of Morphology and Cellular Organization

Our next question was whether distinct cell morphology and organizational patterns were associated with functional differences in phenotype. We started by investigating invasion and colony formation, key properties important for tumor growth and metastatic dissemination.

Using our cohort of cell lines, we performed arrayed colony formation assays (**Figure S8A,B**) and Boyden chamber invasion assays (**Figure S8C-D**) on each of the cell lines. We measured invasiveness and clonogenicity associated with distinct morphology and organizational patterns. While our measurements were limited by sample size and high intra-group variance, we noted that in general, polygonal and aggregated cells had both lower colony formation proficiency (colony count, occupied plate area) and transwell invasiveness compared to cells with other morphologies and organizational patterns. Additionally, all high outlier measurements in the colony formation area measurement were irregular or spheroid cell lines, which typically fall into the dispersed and multilayered organizational categories. KP-4, Suit2, Patu8988T, and MIA PaCa2 featuring irregular/spheroid morphology and dispersed/multi-layered organization were the cell lines with the highest colony formation proficiency while Panc1, KP-4, and CFPAC-1 all representing irregular and dispersed cell lines were the most invasive cell lines. These measurements of invasiveness and colony formation were generally correlated: the Pearson correlation coefficient of log transformed colony formation assay measurements was 0.845 (*p* = 7.2e-5) between colony count and colony area covered, 0.561 (*p* = 0.030) between invasion area and colony area, and 0.408 (*p* = 0.13) between invasion area and colony count.

A subset of the PDAC cell lines in DepMap (n = 30) have also been profiled in a metastatic tropism mouse xenograft model where barcoded cell lines were pooled and delivered via intracardiac injection (*63*). Using this model, we assessed the predictive power of cellular morphology and organizational pattern to stratify metastatic proclivity. We found that consistent with our *in vitro* data (**Figure S8A-D**), irregular/spheroid morphology and multilayered organization were associated with numerically higher rates of metastasis (**Figure S8E,F**). Of the multilayered cell lines, Suit2, Psn1, and MIA PaCa2 have each been independently reported to have high metastatic capacity in various *in vivo* metastasis models (*64*, *65*). Additionally, human pancreatic cancer cells with an ameboid phenotype were recently found to be more proficient at perineural invasion relative to an isogenic line with epithelial morphology (*66*). It is possible that spheroid and multilayered lines may be better suited for anchorage independent growth, a key property enabling seeding at distant metastatic sites (*67*). However, we caution that the intracardiac injection model does not recapitulate locoregional mechanisms of metastasis, such as PDAC metastasis to the liver (*68*). Additionally, variability has been reported in the relative metastatic proclivity of the PDAC cell line models described here (*69*). Nevertheless, our collective data suggests that the mechanisms underlying morphological and organizational patterns may also regulate metastatic potential; however, additional validation with larger cohorts and improved preclinical metastasis models is required to fully dissect this phenotypic association.

### Mapping Tissue Structure and Transcriptional State

Finally, to augment the clinical relevance of our approach to studying associations between morphology and cell state, we explored the relevance of cell morphology in the tissue context. PDAC has previously been described by two predominant malignant cell subtypes, namely the classical and basal-like, which display differences in chemosensitivity, invasiveness, and prognosis (*70*). It is known that 3D organoids have a more classical transcriptional state while 2D cell lines skew to a more basal-like or mesenchymal cell state (*71*). We confirmed that this is true even stratified by our morphological subtypes (**Figure 5A,B**). Given that organoids typically adopt a glandular morphology with a central lumen similar to classical glands in human PDAC specimens while 2D cell lines exhibit mesenchymal sheet-like arrangements and are unable to form glandular structures, we hypothesized that morphological heterogeneity in human tumor specimens may also correlate with transcriptional state.

**Figure 5.**
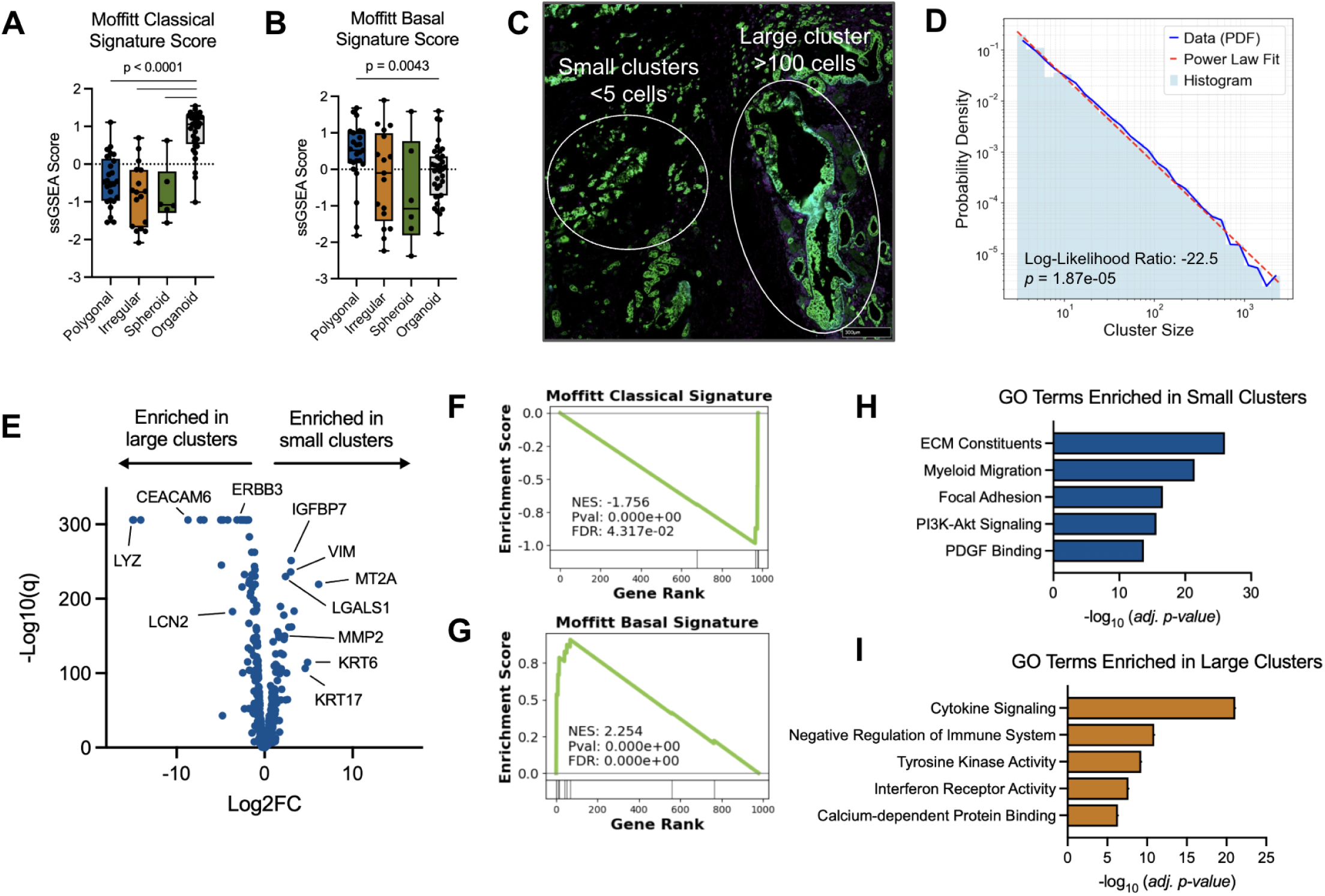
*In situ* tissue morphology in human PDAC correlates with transcriptional subtype. (A-B) Classical (A) and basal-like (B) ssGSEA signature scores (*78*) in 2D cell line models stratified by morphology and in 3D human PDAC organoid lines. (C) Representative comparison of small clusters versus large clusters in DE analysis visualized using immunofluorescence of PanCK (green) from a patient-derived tumor tissue section. (D) Histogram of cluster size distribution plotted on log-log axis. Solid blue line depicts the empirical probability density function, and the dotted red line depicts a power law fitted line. (E) Volcano plot depicting differentially expressed genes in small clusters (right) vs large clusters (left). (F-G) GSEA enrichment plot of classical (F) and basal-like (G) transcriptional subtype signatures (*78*) in small clusters. (H-I) Barplot of top GO terms enriched in small clusters (H) and large clusters (I).

Using a dataset of human PDAC specimens profiled with SMI at subcellular resolution using a 990-plex mRNA panel (*72*), we assigned cancer cells into clusters using a spatial clustering algorithm (DBSCAN; **Methods**). Interestingly, cluster sizes followed a power law distribution (likelihood ratio test, *p* = 1.87e-05), and accordingly we classified individual cells according to their membership in small clusters of 5 cells or less versus cells in clusters of 100 cells or more (**Figure 5C,D**). We performed differential expression analysis and visualized differentially expressed genes in small versus large clusters (**Figure 5E**). Strikingly, the most differentially expressed genes in the small clusters were mesenchymal genes including *VIM*, *LGALS1*, *KRT6*, *MMP2*, and *KRT17*, while the most differentially expressed genes in large clusters were genes associated with the classical state including *CEACAM6*, *LYZ*, *EPCAM*, *AGR2*, and *ERBB3*.

We performed gene set enrichment analysis on the top overrepresented genes in the small and large clusters (**Methods**). Small clusters had enrichment of gene sets related to ECM constituents (*p_adj._*= 9.10e-27), myeloid migration (*p_adj._*= 3.70e-22), focal adhesion (*p_adj._* = 2.45e-17), PI3K-Akt signaling (*p_adj._* = 2.32e-16) and PDGF signaling (*p_adj._*= 1.88e-14). Large clusters featured enrichment of terms related to cytokine signaling (*p_adj._* = 7.90e-22), interferon receptor activity (*p_adj._* = 1.25e-10), and RTK pathways (*p_adj._* = 5.59e-10) (**Figure 5H,I**).

## Discussion

In this study, we systematically evaluated the significance of cell morphology and organization in patient-derived PDAC cell lines and tissue specimens. Our work not only provides a comprehensive assessment of morphological heterogeneity, but also a framework, Spatial Morphology and RNA Transcript (SMART), for understanding and classifying the diversity of human *in vitro* cell lines extensively used by the scientific community. We demonstrated that patterns readily observable in cell culture correlate with functional traits including stemness, invasion, and metastasis, as well as molecular properties such as pathway dependency and gene set expression. Furthermore, we developed a new assay to integrate morphologic and transcriptomic measurements in pools of cell lines. We extended our findings to patient tissues to find distinct transcriptional state associations with cancer cell morphology and organization *in situ*. Diverse treatment emergent morphologies were shown to arise in response to both new and established clinical compounds, suggesting distinct mechanisms of acute treatment response. Using our SMART assay, we identified distinct transcriptional and morphological phenotypes associated with treatment using *KRAS* inhibitors. Our findings support known mechanisms of treatment resistance, involving focal adhesion, integrin, and EMT signaling pathways, while also nominating new hypotheses related to interferon signaling. In response to chemotherapy (e.g. 5-FU and gemcitabine), we observed a consistent induction of a morphologic state characterized by increased eccentricity and cell area.

Our work highlights new directions to expand the characterization of morphology as a resurgent surrogate biomarker of cancer cell state. Additionally, the functional and morphological differences of the Patu8988S/T cell lines derived from the same patient tumor demonstrate that intratumoral heterogeneity can be modeled *ex vivo*. While groups like the JUMP Cell Painting Consortium have pioneered the study of many perturbations including small molecules and under- or over-expression of genes in a few model cell lines, we took the complementary approach of profiling a diverse set of cell line backgrounds with a carefully selected set of clinically relevant perturbations. Our approach shares some concepts with the recently reported STAMP technology (*73*), but is unique in its integrated characterization of morphology and cellular organization, and application to large pools of cancer cell lines. Using barcoded pools of cell line models and imaging-based barcode identification, one could envision characterizing cell morphology changes in a diverse set of genetic backgrounds at large scale beyond what can be accomplished in a single academic lab. Recently, technological improvements such as lab automation and microfluidic imaging have also greatly expanded the scale of morphological profiling studies. For example, Recursion Pharmaceuticals has built screening platforms to assay whole genome CRISPR knockout and activation libraries, and large compound libraries to find compounds that can phenocopy genomic perturbations (*74*). Single cell flow cytometry based morphology profiling (DeepCell) is able to perform high throughput image-based cell sorting (*75*), which has proved useful as a screening endpoint to accelerate functional profiling.

Image-based profiling is a relatively low-cost measurement and thus can be a resourceful way of adding complementary phenotypic information to already acquired transcriptomic databases of cell line models such as the Cancer Dependency Map. Expanding imaging measurements to include cancer types beyond PDAC would improve our ability to correlate morphologies with relevant -omic and functional features. Certain features such as multi-layered organization and spheroid or irregular morphology that we identified to be associated with greater invasive capacity should be validated across multiple tumor lineages and experimental settings. Additionally, more widespread measurement of morphology could improve identification of molecular features and distinct dependencies of different morphologies. While GPX4 dependency has been described in the literature as a feature of mesenchymal cells (*49*), it was identified by ssGSEA using RNA expression instead of morphologic characteristics. In our data, we identified several other mesenchymal, or irregular morphology, specific dependencies including *TRA2B,* which was not previously described. Functional and molecular characterization of the role of the *TRA2B* splicing factor is warranted to assess its potential as a cell-state specific therapeutic target.

We recognize several important limitations of our work. Cell lines undergo transcriptional and morphologic drift in culture. While live-cell imaging with holotomography can resolve dynamic changes in morphology, the SMI assay utilizes a static measurement of spatially-resolved cellular transcriptomic state. Our pooled screening approach has several limitations including the lack of barcodes in cell lines to confirm identity, and the limited 1000-plex probe set used to distinguish cells. The use of spatial molecular imaging panels with increased molecular plex such as 6000-plex and whole transcriptome panels that have recently been developed (*76*) could further improve cell line annotation. While we have demonstrated important clinicopathologic ramifications of cellular and tissue morphology and organization in PDAC, validating our findings in larger cohorts and expanding the application of our method to other cancer types is warranted.

In conclusion, our approach highlights the exciting possibility of leveraging integrated morphologic, transcriptional, and functional measurements on multiple patient avatars in response to drug treatment to comprehensively model patient disease in real-time (*77*).

## Materials and Methods

### SMART Framework

We derived cell morphology data from holotomography, CellPainting, and spatial molecular imaging-based measurements, each providing a distinct advantage. Holotomography provided ultrahigh resolution dynamic live cell imaging, while CellPainting provided higher throughput characterization and quantification of structural features such as actin fibers. Finally, spatial molecular imaging enables high-plex transcriptomic feature assignment to static cells in a single pooled assay.

### Holotomography

We performed holotomography on cultures of human PDAC cell lines under KRAS inhibitor (RMC-6236 or MRTX1133) treated and untreated conditions. Briefly, 300,000 cells were plated on Ibidi 35 mm culture dishes with uncoated polymer coverslips (81156) or 25 x 75 mm microscope cover glasses (Mercedes Scientific, MERR2575) with Ibidi removable culture chambers (80841) and allowed to adhere overnight. The next day, cells were placed on the Nanolive 3D Cell Explorer 96focus for a 48- or 72-hour acquisition.

### CellPainting Assay

Cells were stained with fluorophore conjugated phalloidin (Thermo Scientific; A12380), fluorophore conjugated concanavalin A (Thermo Scientific; C11252), and DAPI and imaged at 10x or 20x using the Nikon AXR. For certain experiments, we switched out the concanavalin A conjugate stain for an anti-paxillin primary unconjugated antibody and subsequent staining with secondary antibody. CellProfiler quantification only utilized the phalloidin stain and the DAPI nuclear stain. Imaging batches were collected for all cell lines and treatment conditions in the same imaging session with identical microscope settings.

### Morphology Quantification with CellProfiler

Raw .tiff images corresponding to each channel (DAPI, phalloidin, and concanavalin A) were processed using CellProfiler version 4.2.6. We designed a pipeline to quantify image features by first extracting primary objects using the ‘IdentifyPrimaryObjects’ module and the DAPI channel image as input with typical pixel diameter range of 20-70 units, global thresholding with minimum cross-entropy, a threshold smoothing scale of 1.3488, correction factor of 1.0, and used intensity to distinguish clumped objects. We then used the ‘IdentifySecondaryObjects’ module to identify cell bodies propagated from the identified primary objects and the actin channel stain. Here, we also used the global minimum cross-entropy thresholding method with a regularization factor of 0.005. Then, we used the ‘MeasureObjectIntensity’, ‘MeasureObjectSizeShape’, ‘MeasureObjectNeighbors’ to quantify cellular characteristics and exported using ‘ExportToSpreadsheet’.

### Spatial Molecular Imaging

We performed spatial molecular imaging (CosMx, Bruker/Nanostring) on treated and untreated cultured cells. For monoculture experiments on Aspc1 cells, we treated cells with 1 µM MRTX1133, and for experiments performed on the pooled cohort of cell lines, we treated cells with 100 nM RMC-6236. This is due to MRTX1133 being an allele specific KRAS G12D inhibitor, and RMC-6236 providing inhibition across the broader set of KRAS alleles present in our cohort. Briefly, we seeded cells in various configurations (either a single chamber per slide or multiple smaller Ibidi chambers per slide) at a density of 1,000 cells per 1 mm^2^ on glass SuperFrost plus slides (Fisher 12-550-15) that were precoated overnight with 0.1mg/mL Poly-D-lysine solution (Gibco A3890401). For pooled experiments, cells were seeded as equi-cellular mixtures. Cells were allowed to adhere overnight and were treated the following day with drug compounds (MRTX1133 1µM, RMC-6236 100 nM, 5-FU 3 µM, gemcitabine 100 nM) for 72 hours.

Cells were fixed with 10% NBF at room temperature for 30 minutes, washed 2x in PBS for 5 minutes, and dehydrated with 70% EtOH at 4° C until slide preparation for CosMx SMI. Slides are stable for up to 2 weeks. Slide preparation followed a modified procedure from Bruker MAN-10184-03. Briefly, slides were rehydrated in 5-minute incubations in 70% EtOH and 50% EtOH, washed with PBS, permeabilized with PBS-T for 10 minutes, and washed twice more with PBS. Samples were digested with 5 ug/mL Endoproteinase GluC (NEB P8100S) for 15 minutes at 40 °C in a hybridization chamber. After incubation, slides were transferred to a jar with PBS, and a 0.0003% fiducial concentration was applied in SSC-T buffer for 5 minutes. The slides were washed once in PBS then transferred for post-fixation to 10% NBF for 1 minute, and then two 5-minute incubations with NBF Stop Buffer (Tris-Glycine buffer) before transferring to PBS for 5 minutes. A freshly prepared 100 mM NHS-acetate mixture (ThermoFisher Scientific 26777) was applied to the cells and incubated for 15 minutes at room temperature. Slides were twice transferred to 2x SSC buffer for 5-minute washes and stored at 4 °C prior to in situ hybridization. We utilized the CosMx Human Universal Cell Characterization RNA Panel (Bruker, CMX-H-USCP-1KP-R), targeting 950 human genes, and a 50-target add-on panel set for all our experiments. These probe sets were individually denatured at 95 °C in a preheated thermal cycler for 2 minutes and crash cooled on ice for 1 minute. We added RNase inhibitor, Buffer R, and DEPC-treated water according to suggested ratios and incubated slides in a hybridization tray for 16-18 hours at 37 °C overnight. We then performed 2 stringent washes with prewarmed 50% deionized formamide in 2x SSC for 25 minutes each, followed by a transfer to 2x SSC until staining. We stained with DAPI, CD298/B2M, PanCK, and CD45 according to the recommended concentrations in the slide preparation manual (Bruker, CosMx FFPE Slide Preparation RNA Kit), first for 15 minutes of DAPI, and then 1 hour with the protein cocktail. We washed slides 3x in PBS for 5 minutes each and stored at 4 °C until being run on the instrument. We used configuration D for cell segmentation and configuration A for pre-bleaching.

### Invasion Assays

We performed transwell Boyden invasion assays to assess invasion and migration capabilities in each of our cell lines. Cells were serum starved for 24 hours and then seeded at 50,000 cells per Matrigel coated insert (Corning 354480) with 500 ul serum containing media below the insert and 500 ul serum free media containing cells seeded on top of the insert. Cells were incubated and allowed to invade for 48 hours and stained according to Diff-Quik staining kit instructions. Invasiveness was quantified by taking brightfield images of 5 representative fields of view and quantifying stained areas within the field as well as using the ‘*Analyze Particles*’ module in ImageJ to count cell bodies.

### Colony Formation Assays

Cells were detached using TrypLE and seeded in 24 well plates at a density of 500 cells/well. After 5 days, cells were stained using crystal violet staining solution composed of 0.125 grams of crystal violet in 50 milliliters of 20% methanol. Colonies were imaged against a bright white background and quantified using the ‘*Analyze Particles*’ module in ImageJ to measure average colony diameter, covered plate area, and total colony count within each well.

### Cancer Dependency Map Analysis

Genome-wide CRISPR dependency effect sizes were obtained from the DepMap 24Q2 Chronos dataset. These data are publicly available through the DepMap portal. These dependency effect sizes are quantified as Chronos scores, a statistical measure inferred from a population dynamics model that corrects for sgRNA efficacy, screen quality and cell intrinsic growth rates, and bias related to DNA cutting toxicity. Dependency scores are continuous values that range in magnitude but have empirically demonstrated that essential genes commonly have Chronos score < -1, and unexpressed genes commonly have Chronos score equal to 0. RNA expression data were obtained from the DepMap 24Q2 and were quantified as log_2_(TPM+1).

Functional MetMap metastasis data were retrieved from the MetMap 500 dataset (*63*) located online at (https://depmap.org/metmap/data/index.html). This dataset contains metastatic potential and penetrance of 488 barcoded cell lines to 5 target organs (bone, brain, kidney, liver, lung). Metastatic potential is a continuous variable on a log10 scale, ranging from -4 ∼ 4 that quantifies the barcode abundance of cell lines within a tissue. Values <= -4 refer to non-metastatic lines, -4∼-2 are weakly metastatic, but with low confidence, and >= -2 means metastatic with higher confidence (*63*). Penetrance refers to the percentage of animals that the cell lines were detected via barcode sequencing, ranging from 0∼1.

### Cluster Analysis in Human Tissue

We utilized a primary human PDAC dataset composed of 13 resection specimens profiled with 990-plex spatial molecular RNA imaging as described previously (*72*). We first classified annotated cancer cells into large or small clusters using DBSCAN, a spatial clustering algorithm. Within each field of view, we applied the DBSCAN algorithm with the eps parameter set to 120 and min_samples to 3. Clusters with more than 100 cells were classified as large clusters and clusters of less than 5 cells were classified as small clusters. After cluster assignment we performed differential expression using a Mann-Whitney unpaired U test to identify differentially expressed genes between cells in small clusters vs large clusters.

### Differential Expression Analysis

A challenge for the 1,000-plex SMI technology is a limited probe set for performing differential expression analyses. The limited coverage interferes with the ability to identify significantly enriched gene sets due to dropout and the assumption by rank based GSEA analyses and commonly used gene sets that the provided gene list represents the entire transcriptome. We therefore supplemented traditional rank based GSEA analysis using predefined gene sets with an overrepresentation analysis method implemented by gProfiler (https://biit.cs.ut.ee/gprofiler/gost). Significantly enriched genes in a condition above a Benjamini Hochberg adjusted p-value threshold of 0.01 and a log2fc difference in means threshold of 0.25 were utilized to identify significantly enriched gene sets. For all gProfiler analysis, we capped the maximum gene set size to 500 genes.

## Acknowledgments

We thank the Koch Institute’s Robert A. Swanson (1969) Biotechnology Center for technical support, specifically the Nanowell Cytometry Platform. We would like to acknowledge Sueda Cetinkaya and Andy Aguirre for assistance obtaining PDAC cell lines. We would also like to acknowledge Justine Johnson, Mary McCarthy, and Shanu Mehta for assistance with Nanolive experiments. Finally, we would like to thank Joshua Choe, Martin Hemberg, Christine Hanko, Jonathan Day, as well as members of the Hwang Lab for helpful discussions; Nicole Lester, Dianne Moschella, Tina Balducci, Misha Pivovarov, Serena Sullivan, Sharon McSorley, Matthew Mues, Anthony Zietman, Daphne Haas-Kogan, Theodore Hong, and Ralph Weissleder for scientific and administrative support.

## Funding

National Science Foundation Graduate Research Fellowship (DG)

National Cancer Institute K08CA270417 (WLH)

Burroughs Wellcome Fund Career Award for Medical Scientists (WLH)

Pancreatic Cancer Action Network Career Development Award (WLH)

## Author contributions

Conceptualization: DG, WLH

Funding acquisition: DG, WLH

Methodology: DG, JMB, WLH

Investigation: DG, RL, YC, MR, JWB

Visualization: DG, WLH

Supervision: WLH

Writing—original draft: DG

Writing—review & editing: DG, WLH

## Data and materials availability

All data are available in the main text or the supplementary materials. Spatial molecular imaging data will be deposited upon publication.

---

**Figure S1.**
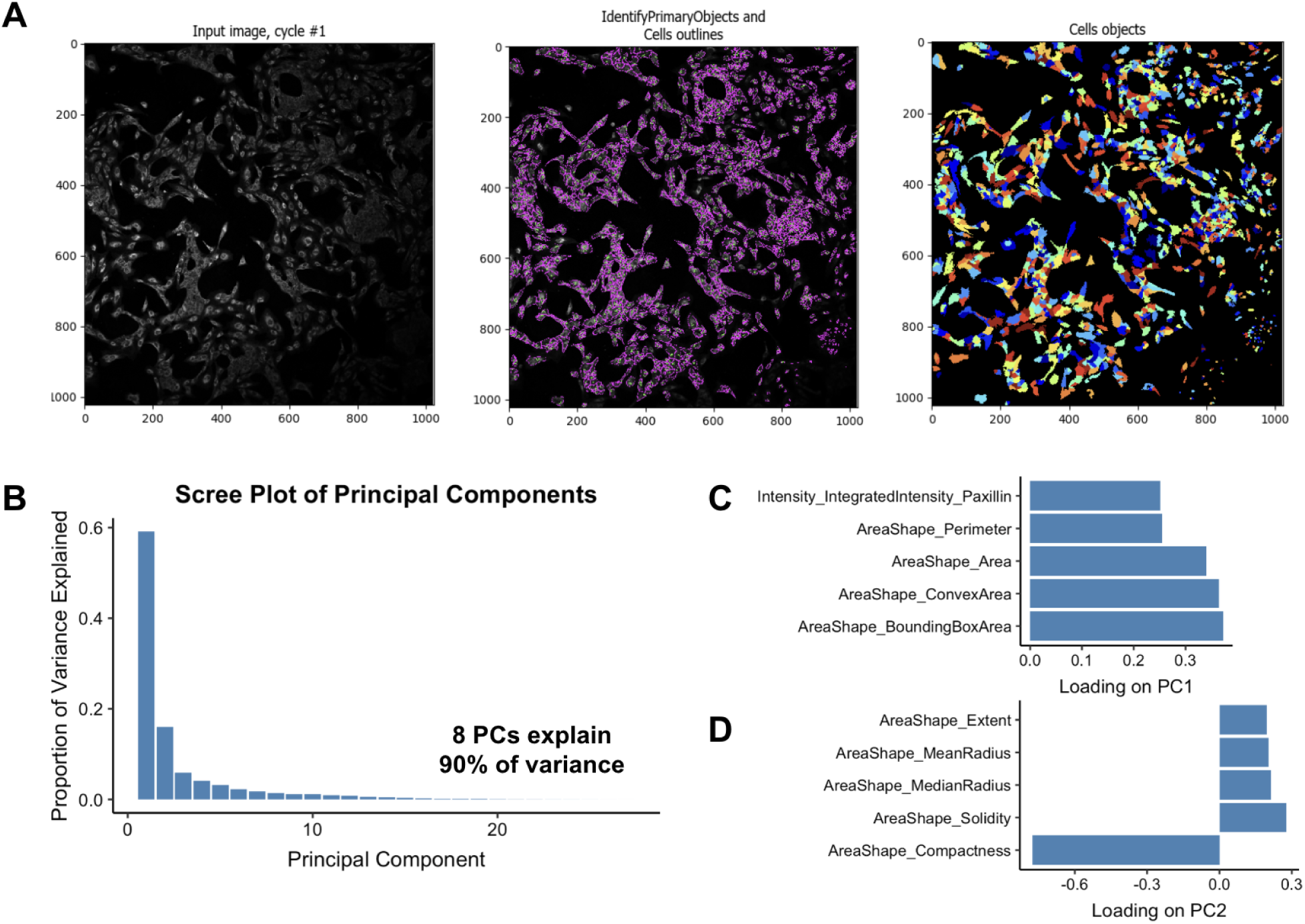
Study schema for arrayed treatment experiment. (A) CellProfiler workflow depicting cell annotation process (**Methods**). *Left*: Merged raw 16-bit input images for DAPI, Phalloidin, and Concanavalin A channels; *Middle*: nuclear and cytoplasm segmentation; *Right*: cell object extraction for downstream measurements. (B) Scree plot of principal components from PCA analysis of morphological features across all profiled cells. (C, D) Barplot of top five features by magnitude in first two principal components.

**Figure S2.**
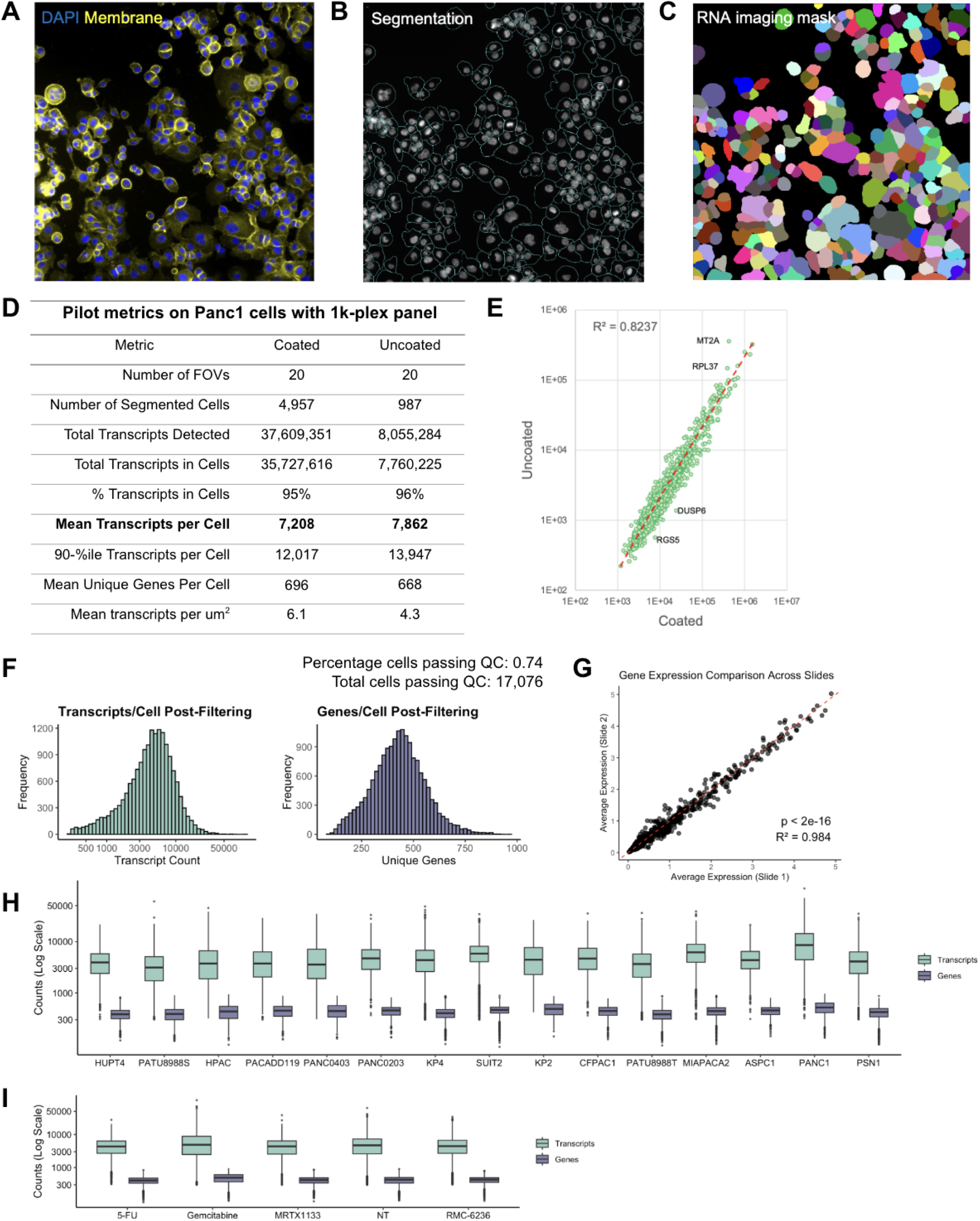
Spatial molecular imaging (SMI) on cultured 2D cells. (A-C) Morphological staining with DAPI (blue) and CD298 (yellow) (A) to aid cell segmentation (B) and creation of RNA imaging masks (C) for transcript assignment and creation of single cell profiles. (D) Metrics from pilot SMI experiments on Panc1 cells using a 1k-plex panel with or without a Poly-L-Lysine slide coating. (E) Scatterplot of transcript abundance from SMI experiments with Panc-1 cells comparing batches of cells prepared with (x-axis) or without (y-axis) Poly-L-Lysine slide coating (**Methods**). (F) Histogram of transcripts per cell (*left*) and unique genes per cell (*right*) in pooled SMI experiment. (G) Correlation of transcript abundance measurements collected from two separate slides of pooled cells. (H) Boxplots of transcript and gene count (y-axis; log scale) for each annotated cell line across all untreated/treated conditions. (I) Boxplots of transcript and gene count (y-axis; log scale) for all cell lines stratified by treatment.

**Figure S3.**
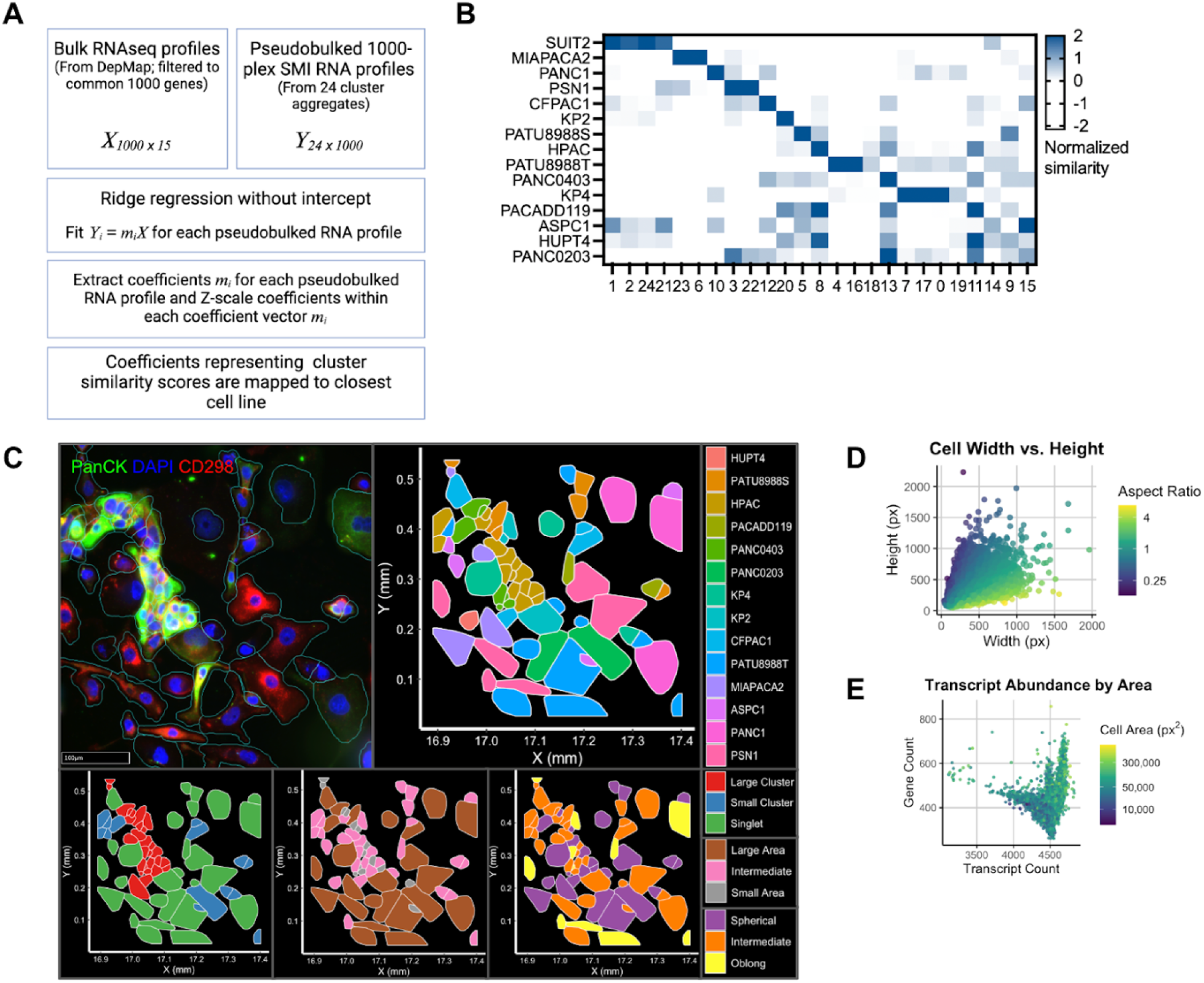
Cell line annotation schema. (A) Flow chart for pooled SMI screen cell line annotation (Methods). (B) Heatmap of Z-normalized similarity scores (color) between each cell line (rows) and cluster (columns). (C) Example slide area with immunofluorescence (top left), RNA imaging mask and cell line assignment (top right), cluster categorization (bottom left), area categorization (bottom middle), and morphologic categorization (bottom right) overlaid. (D) Scatterplot of cell height (pixels; y-axis) versus cell width (pixels; x-axis), colored by aspect ratio. (E) Scatterplot of gene count (y-axis) versus transcript count (x-axis), colored by cell area.

**Figure S4.**
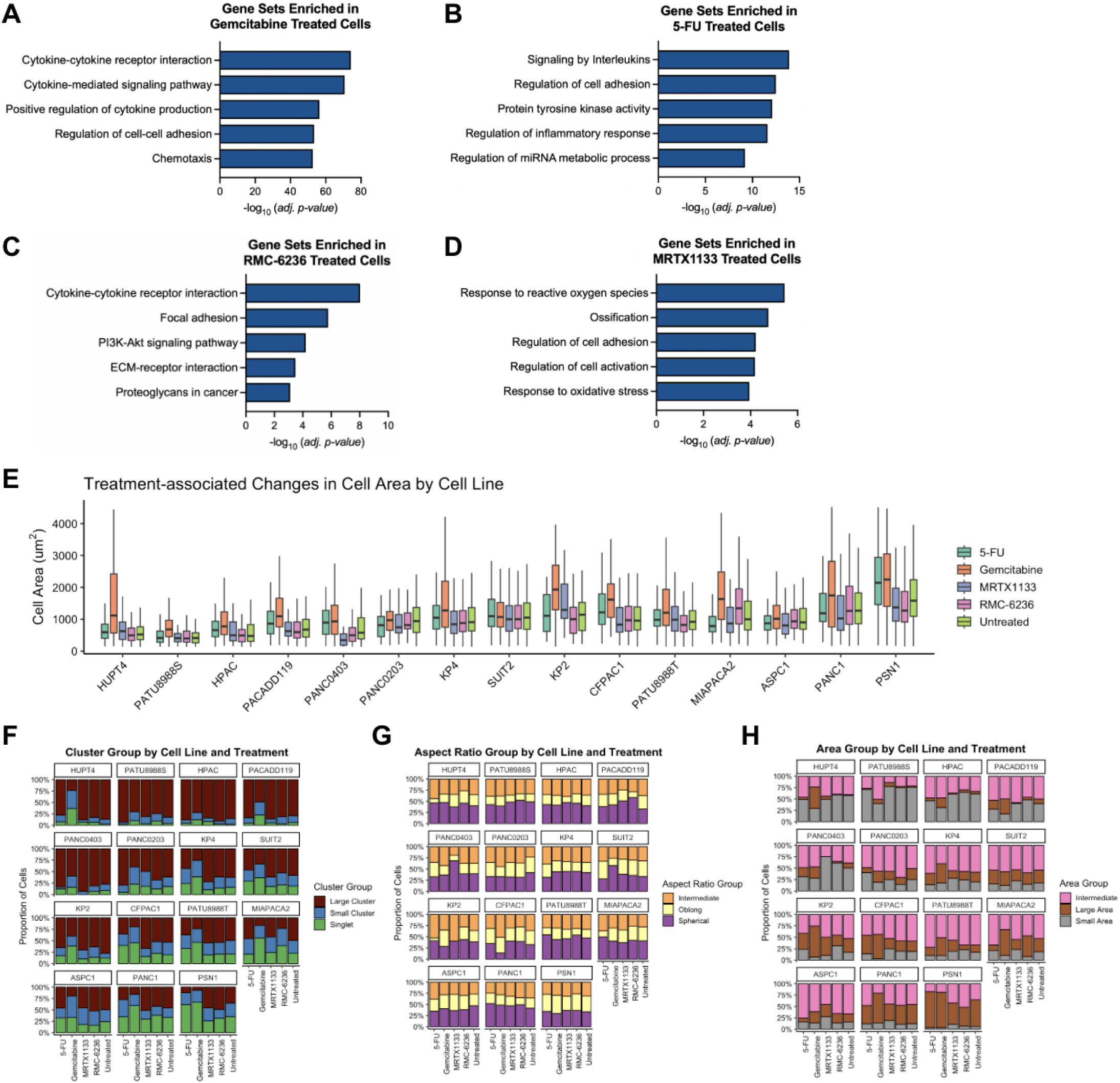
Treatment associated changes in cell phenotype. (A-D) Barplot of top GSEA terms enriched in gemcitabine treated (A), 5-FU treated (B), RMC-6236 treated (C), and MRTX1133 treated (D) versus untreated conditions across all cell lines. (E) Boxplots of cell area measurements for each cell line stratified by treatment condition. (F-H) Stacked bar plots of cell proportions by cluster size (F), aspect ratio (G), and cell area (H) categories stratified by cell line and treatment.

**Figure S5.**
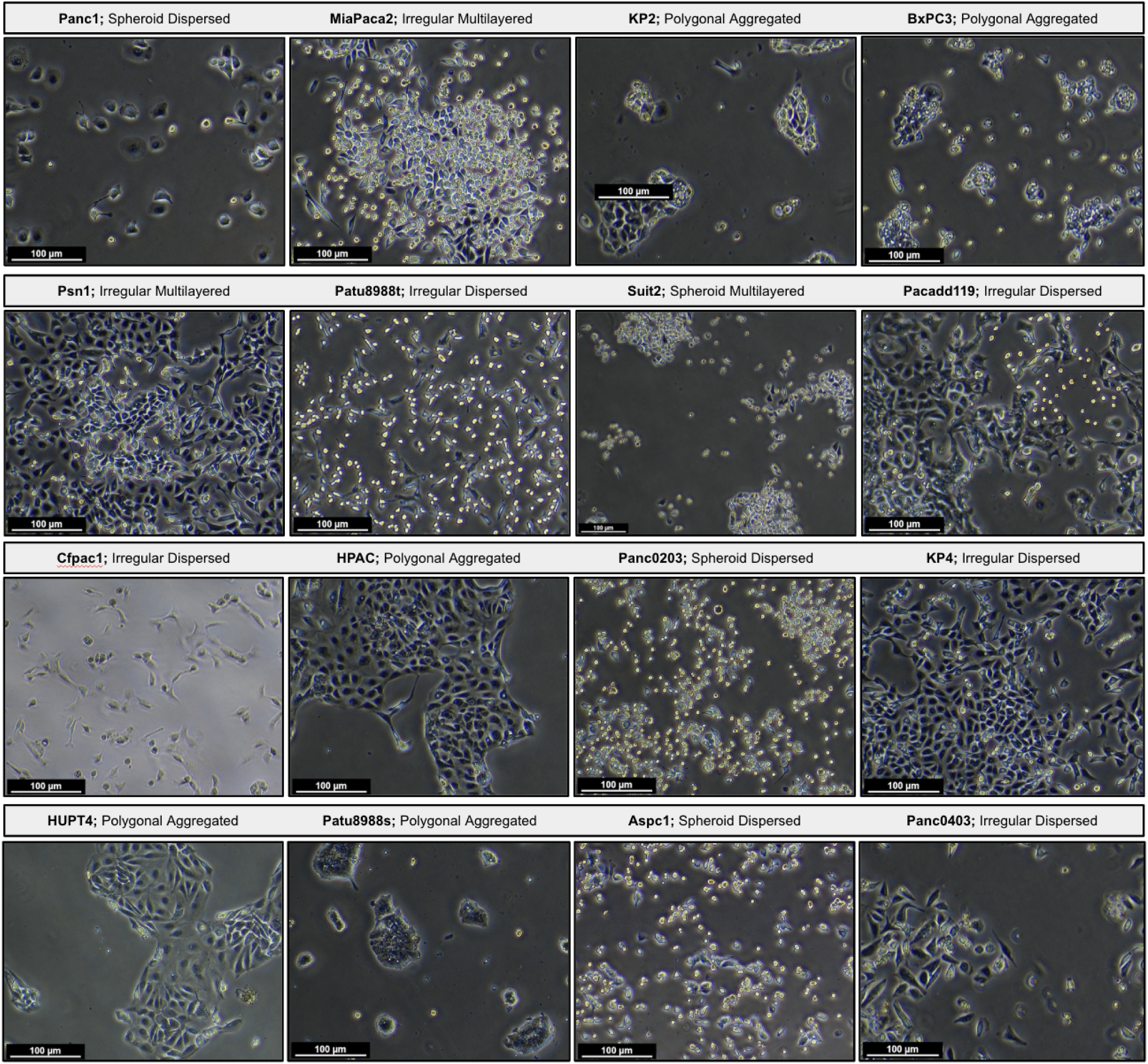
Phase contrast imaging (10x objective) of each cell line included in the custom cohort (*n* = 16) for *in vitro* experiments.

**Figure S6.**
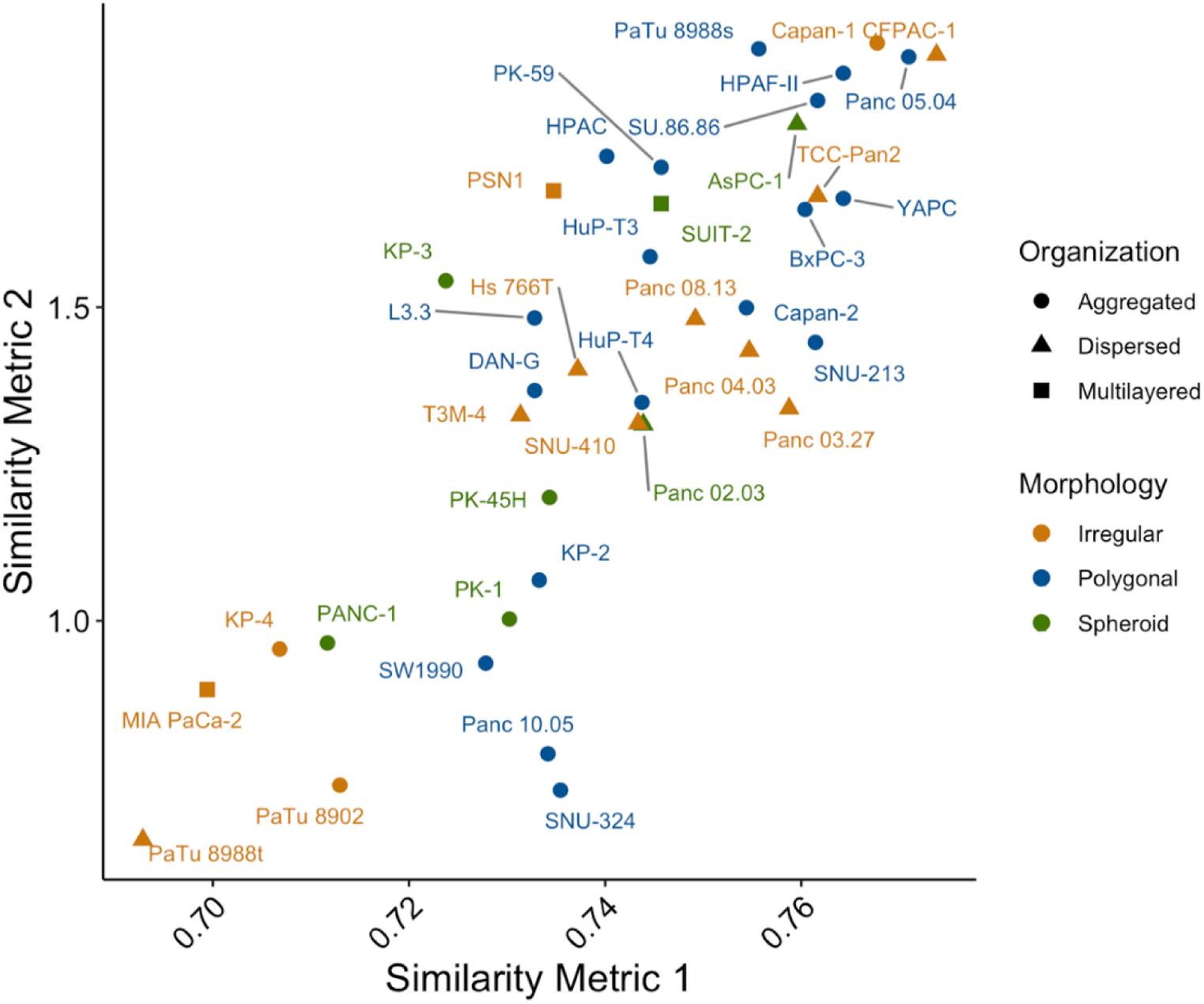
Morphology associated transcriptomes are represented in human tumors. Similarity analysis of cell line transcriptional profile with primary patient tumor RNAseq in the TCGA. Two similarity metrics (Spearman *p* (x-axis) and NES (y-axis), as in Jin, 2023 (*54*)) are used to plot a scatterplot of PDAC cell lines. Dots are colored by morphological categories and have shapes corresponding to organizational categories.

**Figure S7.**
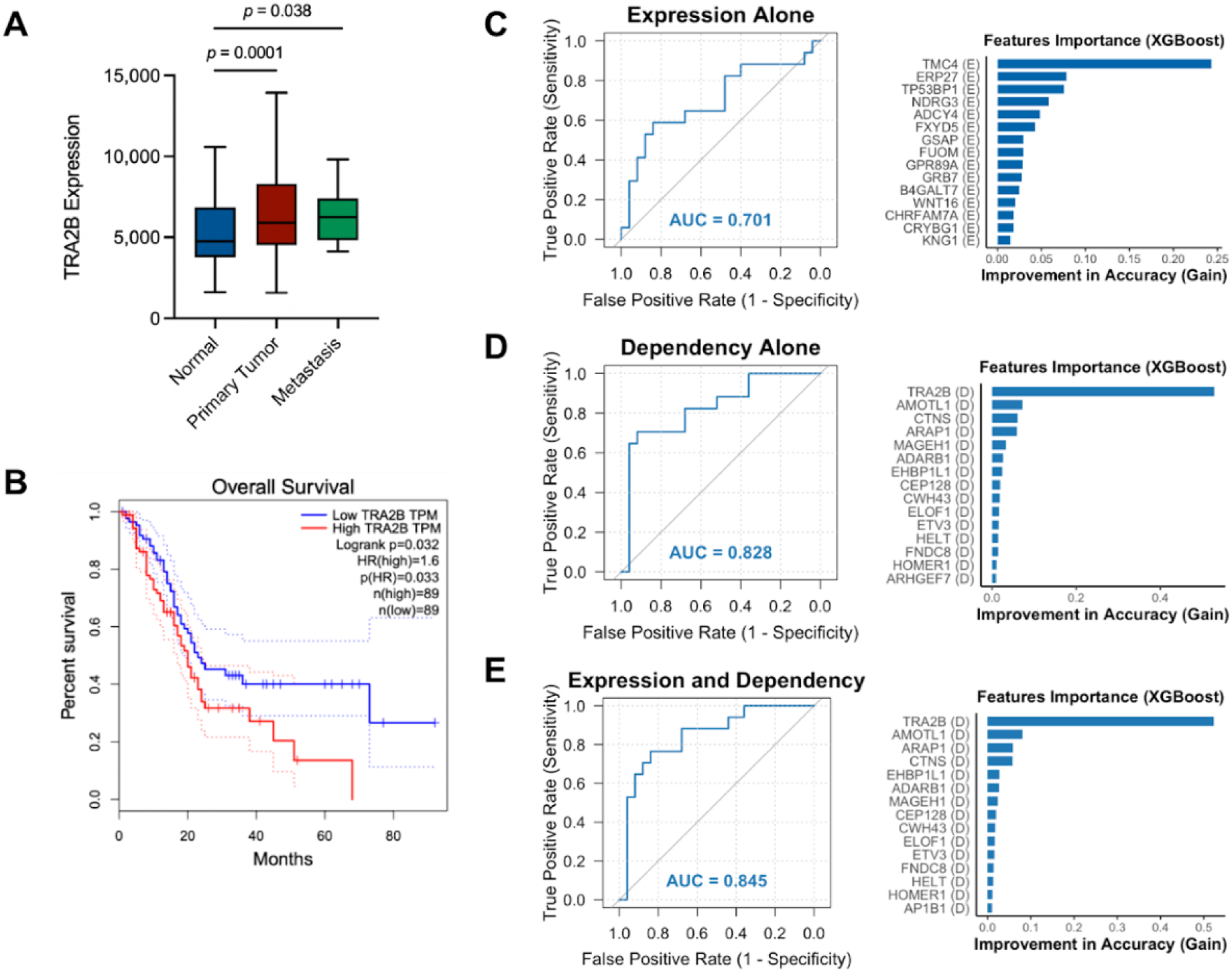
Predicting morphology using expression and dependency measurements. (A) Boxplot of *TRA2B* gene expression in bulk RNAseq data from patient-derived specimens. (B) Kaplan-Meier survival curves of PDAC patients stratified by high or low TRA2B RNA expression based on median expression (GEPIA cohort (*79*)). (C-E) ROC curves and feature importance barplots of XGBoost machine learning models trained on expression and dependency data to predict cell line morphology. AUCs are calculated using leave-one-out (LOO) train test splits and 5 fold cross validation.

**Figure S8.**
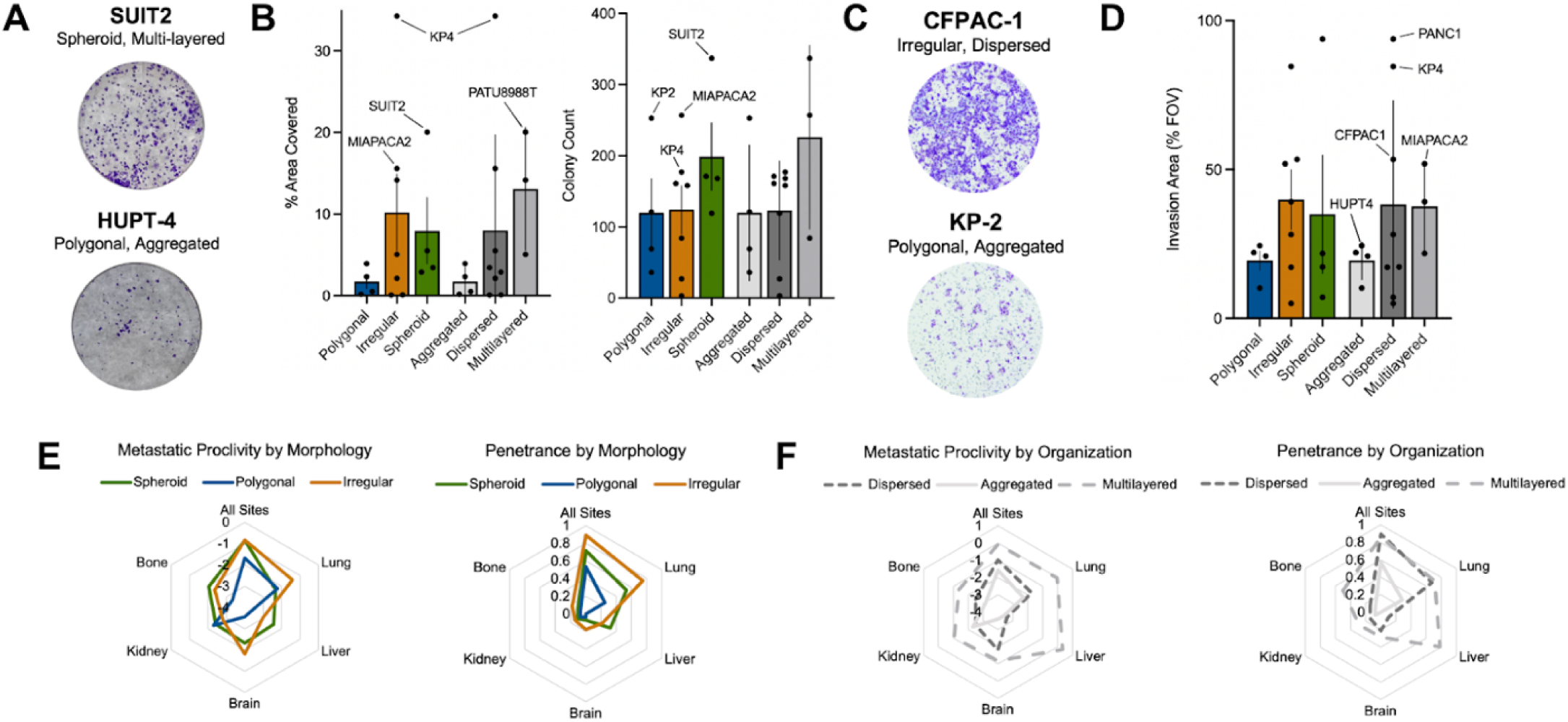
Morphology and cellular organization are associated with metastatic proclivity. (A) Representative image of colony formation assay plate for SUIT2 and HUPT-4 cell lines. (B) Barplots depicting plate area covered (left) and colony count (right) for each cell line model stratified by morphology and organization. Each dot represents a representative replicate from a single cell line and error bars represent standard error or mean (SEM) within each category. (C) Representative image of Boyden chamber invasion assay membrane for CFPAC-1 and KP-2 cell lines. (D) Barplots depicting percentage of field of view invaded for each cell line, stratified by morphology and organization. Each point represents the average across 5 representative fields of view and error bars represent SEM. (E,F) Radar plot depicting metastatic proclivity and penetrance for profiled PDAC cell lines, stratified by morphology (E) and organization (F). Plotted values are calculated as mean across cell lines in the categories from measurements made in Jin 2020 (*63*). Metastatic proclivity is measured by barcode abundance in stated tissue while penetrance refers to the percentage of animals that cell lines were detected in metastases.

